# Composing egocentric and allocentric maps for flexible navigation

**DOI:** 10.1101/2025.01.24.634748

**Authors:** Daniel Shani, Peter Dayan

## Abstract

Egocentric representations of the environment have historically been relegated to being used only for simple forms of spatial behaviour such as stimulus-response learning. However, in the many cases that critical aspects of policies are best defined relative to the self, egocentric representations can be advantageous. Furthermore, there is evidence that forms of egocentric representation might exist in the wider hippocampal formation. Nevertheless, egocentric representations have yet to be fully incorporated as a component of modern navigational methods. Here we investigate egocentric successor representations (SRs) and their combination with allocentric representations. We build a reinforcement learning agent that combines an egocentric SR with a conventional allocentric SR to navigate complex 2D environments. We demonstrate that the agent learns generalisable egocentric and allocentric value functions which, even when only additively composed, allow it to learn policies efficiently and to adapt to new environments quickly. Our work shows the benefit for the hippocampal formation to capture egocentric, as well as allocentric, relational structure – and we link the egocentric SR to findings in the lateral entorhinal cortex. We offer a new perspective on how cognitive maps could usefully be composed from multiple simple maps representing associations between state features defined in different reference frames.

## 1 Introduction

One of the most important dichotomies in spatial understanding is that between allocentric versus egocentric representations. Allocentric representations are tied to a reference frame in the outside world, as if there was a form of (at least contextually local) compass. By contrast, egocentric representations are tied to one of a collection of personal reference frames. In work linking spatial representations to spatial behavior (Geerts et al. 2020; McDonald and White 1994), the underlying difference has historically been very closely tied to that between map-based strategies (Tolman 1948) versus taxon-like habits, built out of motor routines and sensory data (O’Keefe and Nadel 1978; Cheng 2012).

There has been a huge wealth of work on allocentricity in spatial processing. Behavioural evidence for the existence of such representations is broad and deep, including a host of experiments that demonstrate the ability of animals to infer shortcuts (Tolman 1948; Chapuis and Varlet 1987). These experiments, as well as others that show that rodents use geometric features of their environment to reorient themselves (Cheng 1986), and evidence of path integration (Müller and Wehner 1988; Regolin, Vallortigara, and Zanforlin 1995), led to the suggestion that a geometric module exists in the brain (Wang and Spelke 2002; Gallistel 1990). Such a module would process the external environment, turning sensory information which is inevitably egocentric into an allocentric form that is then useful, for instance, for navigation (whence allocentric choices have to be converted back into egocentric coordinates to determine movement). However, the notion that this geometric module might play a role in other spatial tasks, such as representing object locations, was not prevalent. Later work, demonstrating the use of external cues for object memory, suggested the possibility of allocentric spatial memory (Burgess, Spiers, and Paleologou 2004). Abstracting away from the domain of space, there has also been work on allocentric views of social hierarchies (Triandis et al. 1985). Equally, most work on predictive representations such as the successor representation (SR; Dayan (1993)) or the default representation (Piray and Daw 2021) has been conducted in allocentric terms.

Allocentric processing is also important for artificial systems. For instance, there is an abundance of work on simultaneous localization and mapping (SLAM; Thrun (2008)), which concerns the problem of building an allocentric representation of the environment while navigating in it. It is again necessary to ground the map in egocentric inputs to infer odometry measurements. Early SLAM paradigms (Smith and Cheeseman 1986) made use of the Extended Kalman Filter (Kalman 1960; Maybeck 1990) to represent uncertain relationships between different features of the environment. Current neuroscientific models of cognitive maps (Whittington et al. 2020; George et al. 2021; Bicanski and Burgess 2018; Chandra et al. 2023) have retained this classical SLAM-like perspective that embeds objects in an allocentric cognitive map, at most using egocentric representations to build such a map which is then used for planning.

Egocentricity has somewhat sporadically been seen as important for navigational processing. Human and animal studies (Burgess 2006; Wang and Spelke 2002) highlight various phenomena implying the use of an egocentric reference frame. A prominent piece of evidence for such an egocentric representation is the presence of alignment effects - in studies where participants had to learn the location of objects from a single perspective and then recognise those configurations from novel perspectives, the recognition time increased linearly in the angular distance between the two viewpoints (Diwadkar and McNamara 1997). Similarly when participants had to point to the imagined relative position of an object from an imagined viewpoint, they responded faster and more precisely when the imagined viewpoint was aligned with the learned viewpoint (Mou et al. 2004). However, current neuroscientific models (Geerts et al. 2020) have equated these egocentric representations with stimulus-response learning, as in a taxon strategy, rather than using them for the sort of planning that is associated with allocentric representations.

Nevertheless, egocentric representations can also be used in planning. One early influential line of work along these lines suggested the notion of so-called deictic representations (Agre and Chapman 1987). These are locally indexed to the agent – a classic example being ‘the object *in my left hand*’ – and have the advantages and disadvantages of automatic generalization (to any other object in the same hand). Deictic representations have attracted some attention in the field of reinforcement learning (Finney et al. 2012; Agre and Chapman 1987). For instance, because of the inherent generalization they afford, they were considered as potential contributors to model-based planning in a partially observable Markov decision process (POMDP). Unfortunately, the results of doing this were rather mixed (Finney et al. 2012) – deictic representations actually worsened learning performance when compared to a fully-propositional representation. The authors attributed this worsening to the history dependence of an optimal deictic policy – the behaviour of the agent is dependent on the task-location of the agent which is only knowable by examining its action history, and this became hard when the history included exploratory actions.

Although not originally couched in exactly deictic terms, a prominent contribution to work on hierarchical reinforcement learning, the Hierarchical Abstraction Machine (HAM) from Parr and Russell (1997), can be interpreted as such. HAMs are typically modest-sized state machines that specify separable and composable parts of a policy. Parr and Russell (1997) considered environments in which there are many, identical, obstacles around which an agent has to navigate on the way to a distant goal. In this case, a single HAM for avoiding an obstacle can be repurposed to avoid the same obstacle elsewhere. If the HAM is specified deictically (‘turn left when the wall on my right disappears’), then it will typically generalize immediately to other instances of the same object even when rotated.

We so far lack a computationally thorough investigation into modern navigational concepts that exploit egocentric representations in environments that might benefit from them. Given the power of allocentrically-based successor representations (SRs), we investigate egocentric SRs and their use in combination with allocentric representations. We build a reinforcement learning agent that combines an egocentric successor representation with a conventional allocentric SR to navigate complex 2D environments. We demonstrate that the agent learns generalisable egocentric and allocentric value functions which, even when only additively composed, help an agent learn policies efficiently and adapt to new environments quickly. Our work shows the benefit for the hippocampal formation to capture egocentric, as well as allocentric, relational structure. We relate the egocentric SR to findings in the lateral entorhinal cortex.

## 2 Methods

### 2.1 Paradigm

The agent has to solve a complex maze task in which the locations and nature of the barriers and the reward change periodically. Each task involves navigating a 20 × 20 grid with coloured walls which contains a number of identical opaque barriers, randomly located and oriented. (Fig. 1A). In each episode, the agent starts from a random location at a random orientation and moves until it finds the reward, using the actions **“go-forward”, “turn-90**^**°**^**-clockwise”, “turn-90**^**°**^**-anticlockwise”** and **“turn-180**^**°**^ **”**. After every 1000 episodes, the walls and barriers re-organise and the reward location changes.

**Figure 1:**
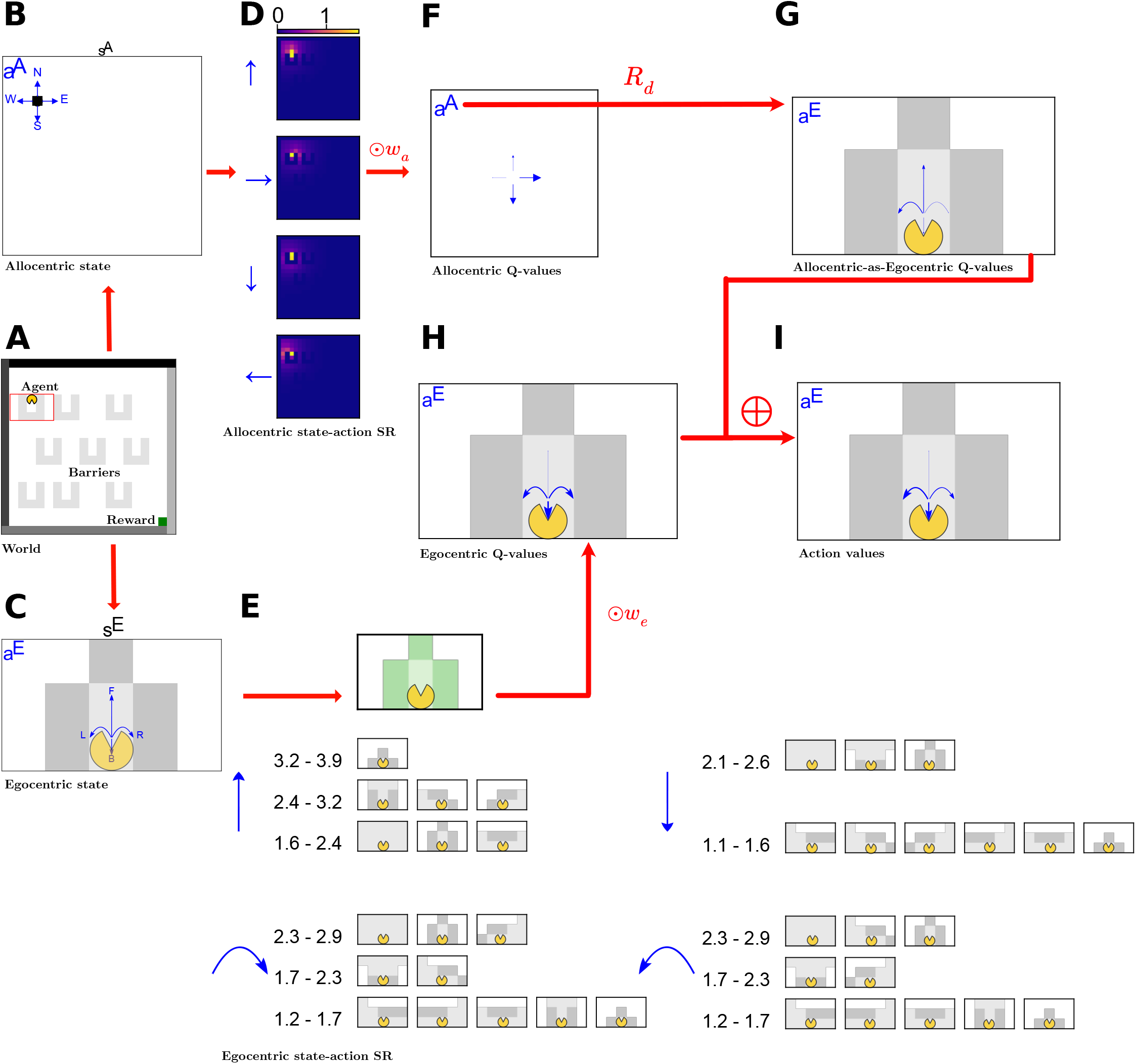
Schematic. **(A)** An example gridworld. The reward is the green box, which is in the bottom right of this gridworld. The grey patches are obstacles to movement and views. In each episode the agent starts at a random location and navigates until it reaches the reward location. The agent is denoted by a yellow pacman figure oriented in the direction it is facing. The red box, along with the orientation of the pacman inside, indicates the egocentric state of the agent. **(B-C)** The agent receives two state representations, corresponding to its allocentric coordinates in the maze **(B)** and its egocentric view, restricted to a local window around the agent oriented in its direction **(C)** and impeded by the barriers. The repeating barriers are in light grey and then there are separate darker shades of grey for each of the four walls. The blue arrows denote the allocentric (**B**) and egocentric (**C**) action sets. **(D-E)** The agent uses the state representations to form state-action successor representations which are used as a basis of its linear function approximation of its action values. **D** shows the allocentric state-action successor representation of the state in **B**. For each allocentric action denoted in blue, the expected future state occupancies are shown as pixel intensities on the 2d grid. **E** shows the corresponding egocentric state-action SR. For each blue egocentric action, the corresponding SR is shown with descending rows denoting decreasing occupancies (in bins, with values shown as numbers on the left). Only the states with the top 60% of occupancies are shown. **(F**,**G**,**H**,**I)** Progressive calculation and combination of action values, indicated by arrow size. **F** shows the allocentric action-values, calculated for allocentric actions, by linear function approximation. Once these are calculated, the allocentric actions are translated, by a direction dependent rotation, into action values for the corresponding egocentric action. These allocentric-state-egocentric-action values are show in **G** and then are combined additively with the egocentric state action-values which are also calculated via linear function approximation **H**, to form our full action-values **I**.

**Figure 2:**
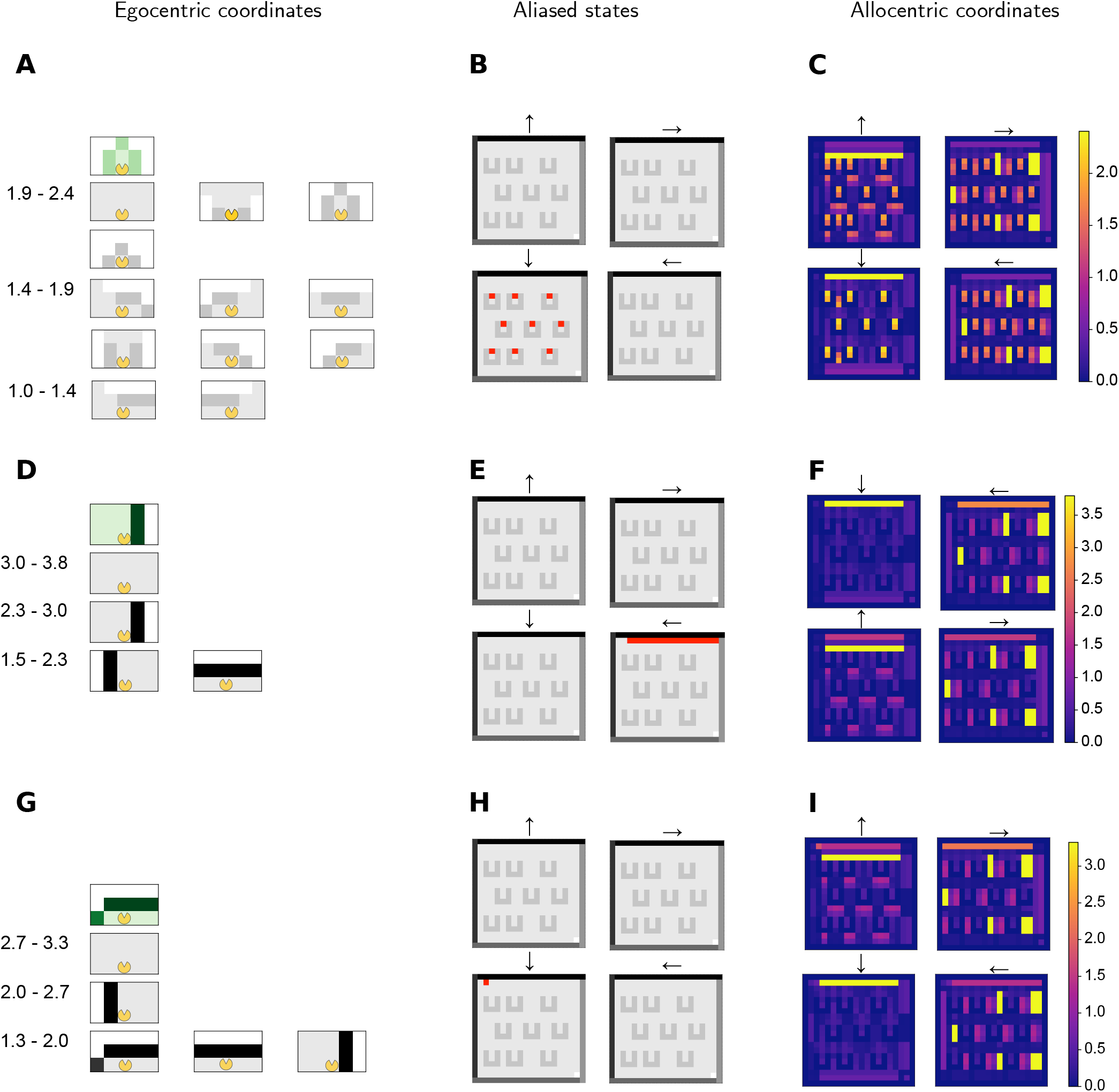
Egocentric Successor Representations. **A-I** Three example egocentric action-independent SRs plotted in both egocentric (A;D;G) and allocentric (C;F;I) reference frames. Note how the SRs look clearly different when visualised in egocentric coordinates but are similar when visualised in allocentric coordinates. The aliased (same egocentric state at different allocentric locations) locations of each egocentric state are shown in B;E;H. The visualisation in egocentric coordinates involves binning the SR vector into 5 equal occupancy bins and then displaying the egocentric states associated with the top 3 bins, in descending rows, so that the highest occupancy egocentric states are displayed in the top row, and so on. For further clarity, we also denote the starting state s in green above the rows. The occupancy bins are shown to the left of the corresponding states. The allocentric visualisation is similar to how we visualised standard allocentric SRs, with pixel intensity corresponding to the expected future occupancy of the state at that location. Since the egocentric state at a certain location is head-direction dependent, we now have four maps, one for each head direction, with the pixel intensity denoting similarly as the discounted expected future occupancy of that egocentric state. While the three SRs are difficult to distinguish when visualised in allocentric coordinates **(C, F, I)**, when visualised in egocentric coordinates they are clearly understandable as the expected future occupancies facing a barrier **(A)**, following a wall **(D)** and facing a corner **(G)**.

At each time-step the agent is supplied with allocentric and egocentric information. The allocentric information (Fig. 1B) is the *s*^*A*^ = (*x, y*) position of the agent in the gridworld. The egocentric information (Fig. 1C), *s*^*E*^, is its immediate visible view, derived from a local window of observations that is centered at the agent and oriented according to the direction the agent is facing. The egocentric view is generically aliased, by which we mean that multiple world states correspond to the same egocentric state, particularly because the barriers are opaque. To calculate the opaque representation, we start with a full egocentric representation centered at the agent, with a fixed horizon of 2. Then we set any pixel values which would be obscured by a block between the pixel and the agent to 0 (the same value as empty space).

### 2.2 Model

We consider a reinforcement learning agent that enjoys a geometric module for converting between egocentric and allocentric coordinate systems, and normally uses state-action SRs following both schemes. At each time-step, the agent receives both allocentric and egocentric state information and uses it to calculate the four separate SRs in each reference frame (Fig 1D, E). These SRs are initialised using trajectories drawn from the uniform policy and are then updated using temporal difference learning as the agent itself learns to behave in the environment. The state-action SRs are used as regressors for a linear function approximation of the state-action value *Q*–function. Here, the allocentric SR is associated with allocentric actions, which are automatically transformed into corresponding egocentric actions using the current head-direction of the agent. The weights of the *Q*–function are learned using *Q*–learning with ADAM (Kingma and Ba 2017).

### 2.3 Action selection

We denote the set of egocentric actions as

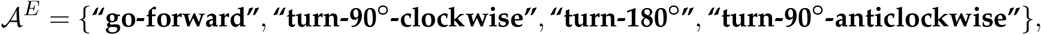

and the set of allocentric actions as

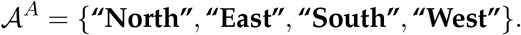

At each time-step, the agent selects an egocentric action *a*^*E*^ ∈ 𝒜^*E*^ by combining information from its egocentric and allocentric SRs. For a given head-direction d we can define a bijection

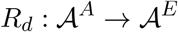

that maps allocentric actions to the egocentric action that would take the agent in that direction. The corresponding egocentric action is *a*^*E*^ = *R*_*d*_(*a*^*A*^).

A consequence of this is that when the allocentric component selects an allocentric action while the agent is facing a different direction, two successive choices will have to be made. Without loss of generality let the allocentric action be **“North”** while the agent faces East. In order to have the agent move North, this action will be transformed into egocentric action **“turn-90**^***o***^**-anticlockwise”** to orient the agent in the North direction, and then the allocentric component would learn that the agent has not moved allocentrically, and would furthermore have to select **“North”** again, which would this time correspond to the egocentric action **“go-forward”**, and now would move the agent to the new allocentric location. To make our model comparisons fair, we only count steps when the agent selects a **“go-forward”** action.

The agent selects a random action uniformly with probability *ϵ* and with probability 1 − *ϵ* selects an action using a softmax policy of the action-values, with softmax temperature parameter *τ*.

### 2.4 Representations

The state-action successor representations are defined in the conventional manner as

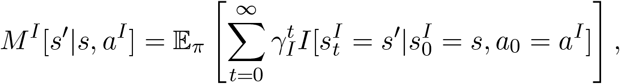

where *γ* ∈ (0, 1) is a discount factor, and *I* is the identity function. These SRs are initialised using transition matrices under the uniform policy and then are updated online using TD-learning with prediction-error modulated learning rate.

The initialisation is according to allocentric and egocentric transition matrices that are created from sample trajectories. Allocentric samples come from following a uniformly random choice of allocentric actions in an empty environment. Egocentric samples come from following a uniformly random choice of egocentric actions in a random environment with barriers.

After observing a transition (*s, a, r, s*′), each state-action SR (*I* ∈{*A, E*}) is updated according to

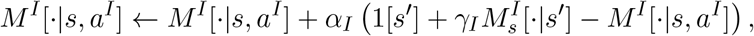

where 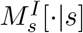, (*I* ∈{*A, E*}) is the action-independent SR, defined as

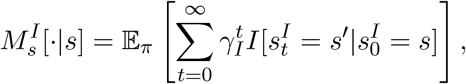

and concurrently learned via the update-rule

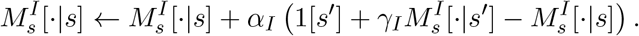

Here *α*_*I*_ is the SR learning rate and 1[*s*′] corresponds to a one-hot encoding of state *s*′.

Note that the allocentric transition graph is larger than the egocentric one, but with sparser connections. Meanwhile the egocentric transition graph has fewer nodes, because of aliasing, but with denser connections. Therefore the SRs have very different magnitudes. This motivated the use of ADAM as a means to adapt the learning rates appropriately for both bases.

### 2.5 Value function

Concurrently, the agent learns a state-action value function using linear function approximation. The agent’s *Q-*values are

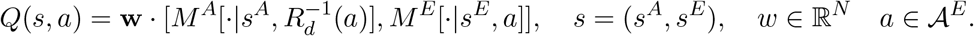

The allocentric SR uses the allocentric action set 𝒜^*A*^ which are then transformed into the corresponding egocentric actions using the head-direction of the agent. The weight vector **w** is learned using Q-learning with ADAM.

All model parameters used for the experiments are shown in Table S1.

### 2.6 Plotting of egocentric SRs

In Figure 3(A-I), three example egocentric SRs are plotted in both egocentric (left column) and allocentric (right column) coordinates. Figure 3B;E;H highlight in red the aliasing of those egocentric states across the first environment. Egocentric coordinates are associated with the (obscured) forward view of the agent. Thus, to show the egocentric SR is to show which particular forward views are prevalent following a given location and head direction, by averaging over future ego-centric ‘pictures’. When plotting in egocentric coordinates, we only display the states in the top 60% of occupancies.

**Figure 3:**
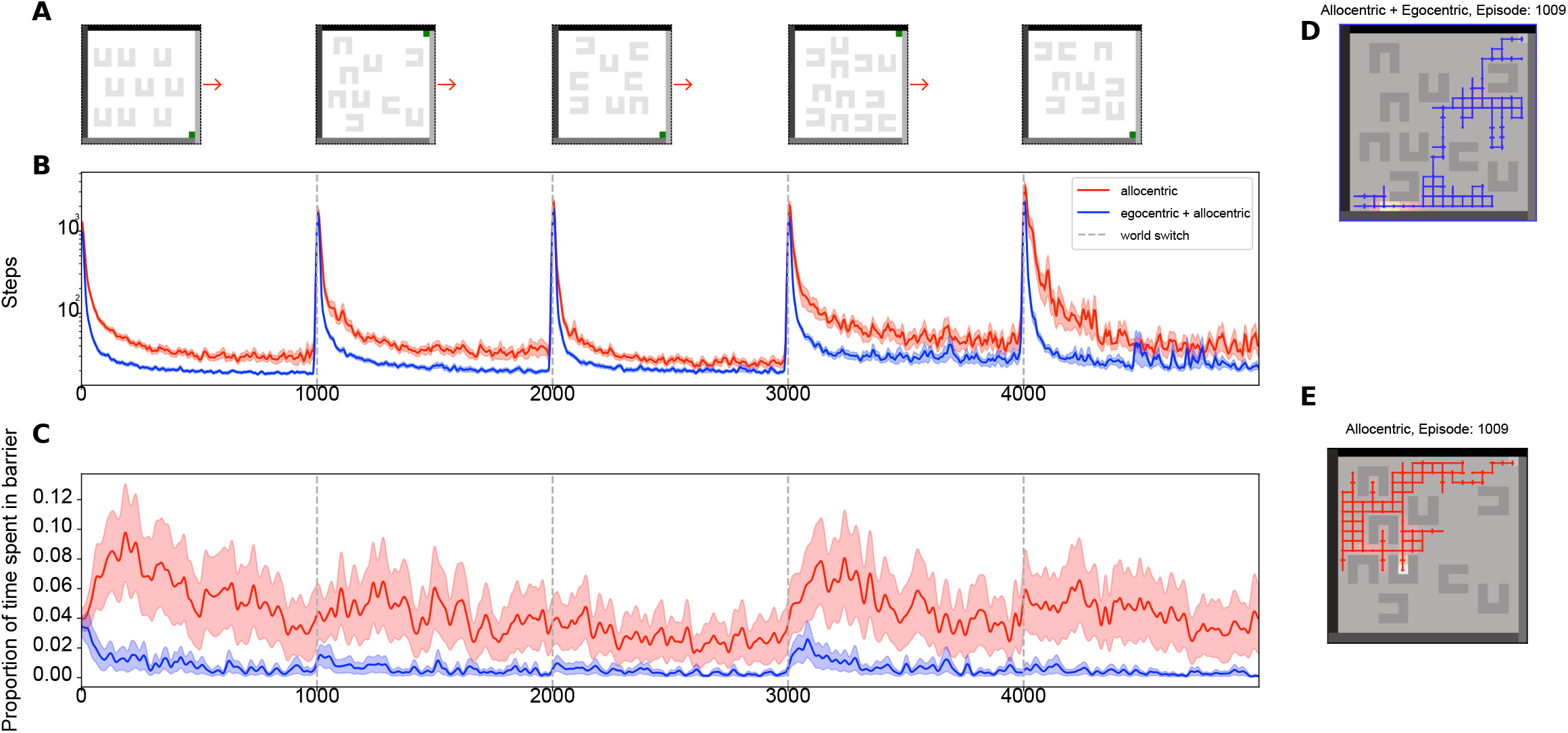
Lesion Comparison - Multi world learning. **(A)** A sample task, in each task we sample 5 random grid arrangements. **(B)** Comparison of two models in the task, averaged over 30 different tasks. One model (blue) is the full allocentric+egocentric agent and the other (red) is a purely allocentric agent, without an egocentric component. The egocentric SR aids the learning of optimal strategies - although both step counts spike at environment switches, the full agent quickly relearns optimal strategies, while the lesioned agent is much slower. **(C)** Plots of the average time spent inside the barriers, for the full and egocentrically lesioned agents, calculated for one task averaged over 30 different seeds of that task. One can see that the full agent has learned to avoid the barriers, while the lesioned agent has not - demonstrating differences in generalisation. **(D, E)** Sample paths taken in the the second environment of a task, soon after an environment switch. **(D)** shows an example path taken by the full agent. It avoids the barriers and gets to the reward quickly. **(E)** shows an example path taken by the lesioned agent at the same point in training. It can be seen to get caught in the new barriers and to avoid areas where there used to be barriers.

The egocentric states for which the SRs are shown correspond to barriers, walls and corners. Note how it might be hard to distinguish C, F and I, but it is easy to distinguish A, D and G. This shows how it difficult to disambiguate these distinct egocentric states by plotting their allocentric frequencies. However, once visualised egocentrically, disambiguation is more straightforward.

### 2.7 Multiple task comparisons

For the multiple task comparison in which we test ablations of the model (Figure 5), we run the agent in a collection of paradigms, each one which randomly samples worlds from different generative models. The generative models either generate barriers with fixed sizes or differing sizes and either fix the orientation of the different barriers or orient each barrier randomly (results shown in Figure 5A-C). We also employ a further generative model that generates fully-random non-compositional gridworlds, using a variant of Prim’s algorithm (results shown in Figure 5D).

## 3 Results

We built an agent which acquires both egocentric and allocentric state-action successor representations, and uses them as the basis for learning *Q*–values to navigate to a goal amidst scattered obstacles. The locations of the goal and the obstacles change every 1000 steps. A sequence of 5 environments is shown in figure 3A.

### 3.1 Egocentric SR facilitates generalisation

The blue line in Figure 3B shows the performance of the agent in this task. The agent rapidly learns to find the goal with efficient paths. Furthermore, having had experience with the barriers, it spends little time being trapped by them (blue line in Figure 3C). The agent has no capacity to ‘learn to learn’, and so had to re-adapt to the change each time in happens (hence the spikes in performance every 1000 trials in Figure 3B). However, the capacity to avoid getting trapped in barriers generalized well after environmental changes (Figure 3C) because of the local consistency of the different worlds in egocentric terms, even though the global structure had changed. This meant that even straight after a change, the barriers could be duly avoided, as shown by the sample path in blue (Figure 3D).

The red curves in figures 3B;C;E result from lesioning the agent to remove the egocentric SR. They show the benefits of having an egocentric representation. The paths are longer (figure 3B), and the obstacles are more of a persistent hazard (figure 3C;E). Forcompleteness, we also plotted the performance of an egocentric-only agent in Figure S2. Furthermore, Figure 3E shows that the lesioned agent avoided empty areas of the world where barriers were previously situated, leading to inefficient strategies. We optimize our lesioned baseline maximally by optimizing the shared hyperparameters between the lesioned and unlesioned agents to maximize lesioned agent performance and then only optimize egocentric hyperparameters on top for the unlesioned agent performance.

### 3.2 The agent learns compositional value functions that facilitate efficient continual learning

To show how this generalisation occurs, we examined the value function learned by the full and lesioned agents. In Figure 4 we plot *V* (*s*) = max_*a*_ *Q*(*s, a*) with *s* = (*x, y, d*) for the full and lesioned agent at the end of learning in the first world, and soon after an environment switch. Due to the linearity of the value function, the separate egocentric and allocentric contributions are easy to compute and are plotted separately. Figure 4B shows the full value function learned by the agent in the first world. Figures 4C;D show how this is constructed from egocentric and allocentric components respectively. Figure 4E shows the difference between the value function when the agent is at a certain direction and the mean directional value function (Figure 4C). For comparison, the value function learned by the lesioned agent is shown in Figure 4F.

**Figure 4:**
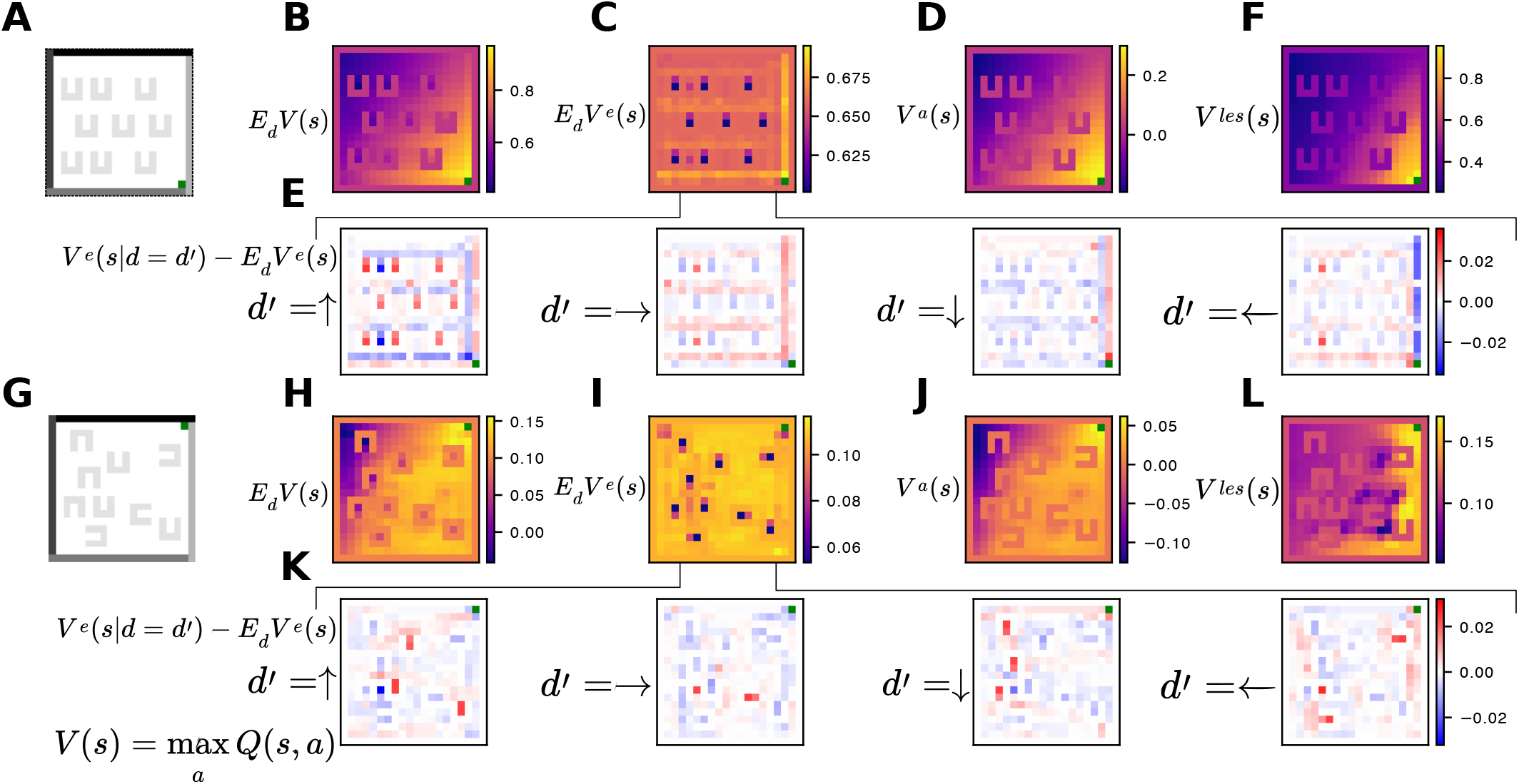
State value functions. **(A-F)** Decomposed state-value function *V* (*s*) = max_*a*_ *Q*(*s, a*) learned by the full and lesioned agents near the end of training in a example first environment, shown in **(A). (B)** shows the full value function learned by the agent, averaged uniformly over head-directions, *E*_*d*_*V* (*s*). **(C, D)** show the egocentric *E*_*d*_*V*^*e*^(*s*) and allocentric *V* ^*a*^(*s*) contributions to this value function. It is a combination between a general value gradient decreasing from the reward location and local perturbations around barriers and walls. We then show in **(E)**, for the unlesioned agent, the direction-based perturbations around the mean egocentric directional value function. This highlights the agent’s preference for wall-following strategies and for avoiding barriers. **(F)** shows the value function learned by the lesioned agent. It learns a similar value function. **(G-L)** Value functions of the full **(H, I, J, K)** and lesioned **(L)** agents, soon after an environment switch. The new environment is shown in **G**. In **(H, I, J, K)** it can be seen that the full agent already has learned the negative value at the new barrier locations and the new reward location (**I, K**), with few artifacts from the previous environment. (**L**) The lesioned agent’s value function: there are clear artifacts from the previous world and it has not learned the negative value of some of the new barrier locations.

Full and lesioned agents learn value functions that solve the environment - learning the reward location and attributing negative value to the barriers. However, in the full agent, the different egocentric and allocentric components represent different aspects of the environment: the allocentric component learns a general value gradient which reflects the reward location but ignores barriers. The egocentric component learns local subtractive value functions that stop the agent from getting caught in the barriers and which generalise across the environment, as well as learning a preference for following the wall that leads to reward.

We then examined the value function of the full and lesioned agents soon after an environment switch. While both agents learned similar value functions for the first world, after 10 episodes in a new environment, the two value functions were substantially different (Figure 4H-L). The lesioned agent had learned the new reward location, but had not learned to associate negative values with the new barrier locations, furthermore there were deleterious artifacts from the previous barrier locations. By contrast, the full egocentric agent had learned to avoid the new barrier locations and did not carry over negative values from previous barrier locations.

In sum, the full value function consists of a combination of the allocentric value gradient and the local egocentric strategies that correct the gradient-following strategy to avoid colliding with the barriers. These local egocentric strategies generalise to other environments that share the same local structure, even though the global structure is vastly different.

### 3.3 Contribution of egocentric component varies with the local consistency in ego-centric terms of the different worlds

In order to investigate the variability in the utility of the egocentric representation across a range of task-types, we re-ran the lesion comparison in tasks with differing levels of local consistency. We compared full and allocentric-only agents in different tasks where the worlds were sampled from generative models that either kept the barriers aligned against reward (Figure 5A;B), or where orientations were randomly chosen per barrier (Figure 5C). We also did this for worlds where the barriers could change their size (Figure 5B;C). We then ran the agent in fully randomly generated worlds where the only restriction was that space had to be fully-connected (Figure 5D).

**Figure 5:**
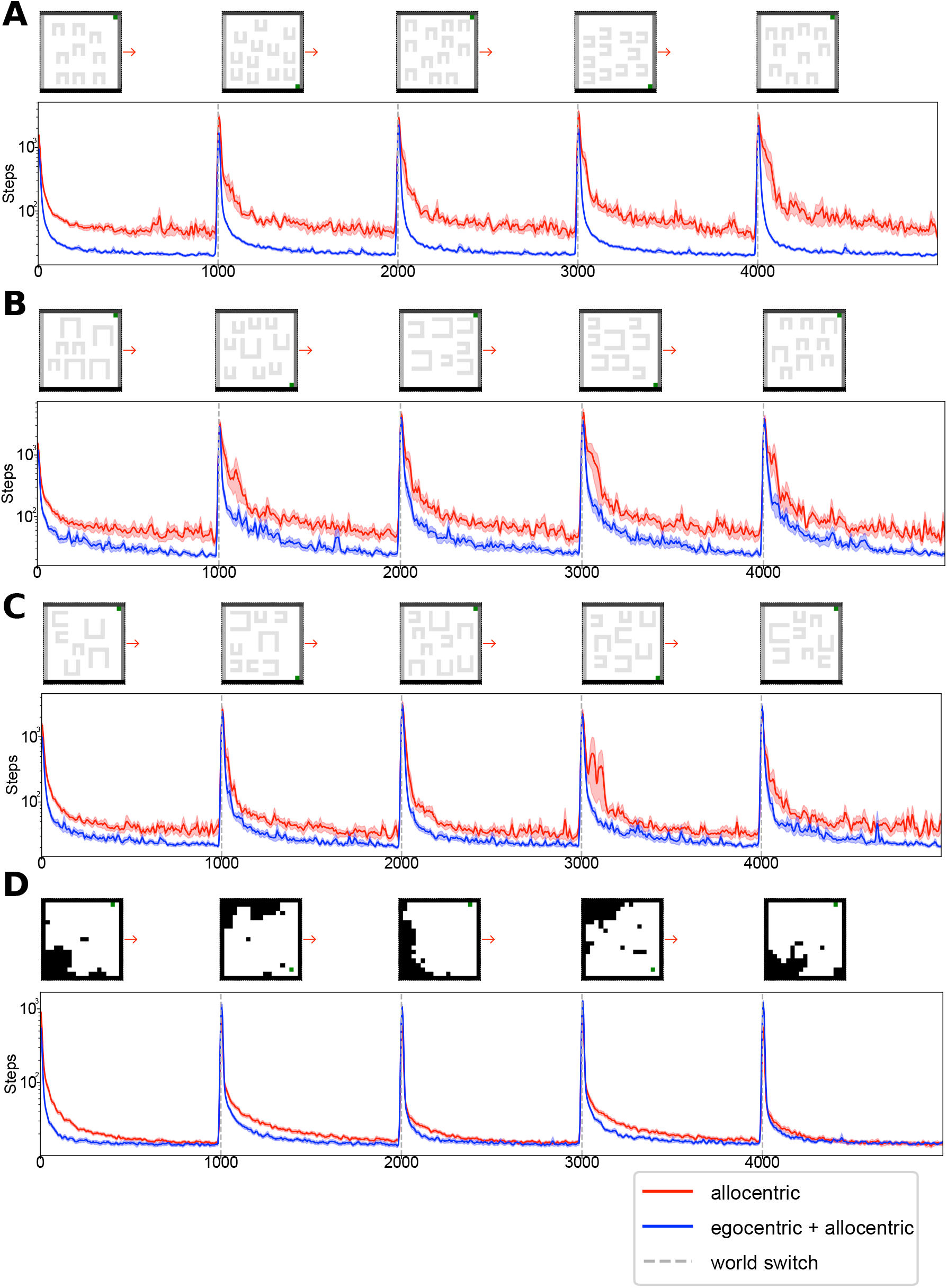
Comparison of the benefits of the egocentric component in other task paradigms. **(A-D)** Comparisons between full (allocentric + egocentric) and lesioned (allocentric-only) agents across paradigms where tasks are being sampled from different generative models. Each plot consists of (on top) an indicative sample task (bottom) the lesion comparison across runs with different tasks drawn from a specified generative model. **(A)** Comparison between full and lesioned agent where environments consist of randomly generated barriers, of the same size, which are all aligned to obstruct reward access. **(B)** Comparison between full and lesioned agent where environments consist of randomly generated different-sized barriers which are all aligned to obstruct reward access. **(C)** Comparison between full and lesioned agent where environments consist of randomly generated different-sized barriers which are all randomly orientated. **(D)** Comparison between full and lesioned agent where environments consist of randomly generated worlds with no shared egocentric structure across environments.

We found a gradient in the differences in performance between the full and lesioned agents that depends on the consistency of local egocentric structure as well as the usefulness of knowledge of such structure. However, inclusion of the egocentric SR is ubiquitously beneficial. This can also be seen by plotting the ratios of the number of steps taken by the unlesioned and lesioned agents, the histograms of which are shown in Figures S3 to S7.

## 4 Discussion

Our results highlight clear benefits of egocentric associative representations in complex spatial processing. We developed a model which combines a standard allocentric SR for global positioning with an egocentric SR which allows it to navigate efficiently and flexibly around multiple complex environments by exploiting their shared local structure. We demonstrated the benefit of such an egocentric SR by comparisons with lesioned models and, thanks to the simple linear structure of the model, unpicked the contributions of the different components to the value function.

Our understanding of the cognitive mechanisms of spatial processing has been dominated by approaches that are focused on allocentric representations. This is prompted by their evident neural signatures in the rodent hippocampal formation including place cells (O’Keefe and Nadel 1978) and grid cells (Hafting et al. 2005). Lesion experiments further corroborate the hippocampal formation’s role in navigation (McDonald and White 1994). Cognitive maps offer a unifying framework for such analyses, with both place cells and grid cells being seen as being key to understanding the neural basis of such a map (O’Keefe and Nadel 1978; McNaughton et al. 2006; Hafting et al. 2005). Cognitive maps were originally introduced as an internal representation of an environment that can be used for flexible behaviour such as finding new routes and paths. More recent work has emphasised the necessity of the maintenance and updating of one’s position in the environment, known as path integration (McNaughton et al. 2006).

Modelling approaches to understanding the role of the hippocampus in spatial processing have been varied, but have concentrated on allocentric representations. For instance, it has been suggested that the hippocampus offers a predictive map of the future (Stachenfeld, Botvinick, and Gershman 2017), viewing place cells as the neural implementation of an allocentric SR and grid cells encoding a low-dimensionality basis set from which to build such a representation. There have also been suggestions that the hippocampal formation performs latent allocentric structure learning on egocentric inputs (George et al. 2021; Whittington et al. 2020) with grid cell and place cell properties being explained by various functions including path-integration (McNaughton et al. 2006; Dorrell et al. 2023; Cueva and Wei 2018) and probabilistic message-passing (Evans and Burgess 2019). Some of these approaches view allocentric representations such as place cells as latent states of a generative model that tries to reconstruct egocentric sensory information (Benna and Fusi 2021; Gornet and Thomson 2024; Spens and Burgess 2024). The cognitive map concept has also been extended as a more general way of organising knowledge to facilitate generalisation and rapid inference (Whittington et al. 2022). By contrast, work on egocentric representations either focuses on translating from egocentric to allocentric representations, possibly a task for retrosplenial cortex (Bicanski and Burgess 2018; Alexander et al. 2023), or, in navigational terms (e.g., Geerts et al. 2020) tends to regard them as being used for simple taxon-based stimulus-response learning rather than the sort of planning and associative structure learning that is equated with allocentric representations. By contrast, our approach, although using RL and dealing with navigation rather than explicit prediction, suggests one should see them in similar terms to the cognitive map formulations described above, and suggests that when egocentric information is informative, we might expect the hippocampal latent code to capture egocentric, as well as allocentric, information.

Egocentric representations were used for planning in Agre and Chapman (1987). Drawing on work on visual routines (Ullman 1984), this made the distinction between capital-P “Planning”, where a smart Planning phase constructs a Plan which is carried out mechanically, and lowercase-p “planning” which is closer to recipes, which they claim is a better description of naturalistic human behaviour. Agre and Chapman (1987) suggested deictic representations, indexical-functional entities such as “the-block-that-I-am-pushing”, as a means to overcome the combinatorial explosion associated with “Planning” using propositions. Deixis has been extensively explored in linguistics (Kaplan and Kaplan 1989), stemming from the introduction of terminologies such as indexicals, which are linguistic expressions whose reference can shift between contexts. The sort of policies learned by the agent in this study include “plans” based around deictic representations such as “the-barrier-that-I-am-facing”, rather than a plan based around an allocentric representation of the form “barrier1” (as distinct from “barrier2”, which might be physically identical, but placed somewhere else in the environment).

Our investigation shows how exploiting compositional representations can benefit performance in dynamic tasks, for instance facilitating continual learning. Our world model is factorized into simple egocentric and allocentric maps, allowing it to represent separately different dimensions of the environment which better generalise to new environments. Indeed, we showed how performing spatial processing in an egocentric reference frame enables a form of passive generalisation across environments which share local structure. This led to fast learning after environmental change. More generally, animals could learn to navigate complex environments by composing simple maps representing different reference frames. These different reference frames do not in general have to be egocentric and allocentric but rather could represent learned priors over other important dimensions that collectively factorise a world model. For instance, evidence about object-based attention (Chen 2012) suggests that objects might provide another useful reference frame that would be closely related to our egocentric suggestion.

Factorised approaches of this sort have been successful in SLAM. The main original SLAM approaches were based on algorithms such as the Extended Kalman Filter, where a robot might navigate by maintaining and updating a mean vector (best estimate) and a covariance matrix (expected error) of the locations of itself and N landmarks. However, the computational complexity of this, mostly stemming from the quadratic nature of the covariance matrix, have more subsequently led to approaches that exploit local connectivity (Lu and Milios 1997; Thrun et al. 2004) or decompose the map into sub-maps (Bailey 2002) rather than learning full global structure. These updated approaches to navigation are conceptually similar to the approach we have advocated, by emphasising the importance of local coordination and representations that are factorised into simpler sub-maps.

Related modeling work on the hippocampus also makes use of compositional representations to facilitate generalisation. Bakermans et al. (2023) employ a meta-learning approach to train a large neural network from pre-determined optimal policies, and thereby show how using state spaces which are composed from reward and object-vectors can help one infer optimal policies in new environments better than using pure spatial representations. Though similar in flavour, there are various differences with their approach, including our application of RL with a simple linear representation to show how a factored representation allows quicker learning and generalization in real-time.

A separate motivation for our work was hierarchical abstract machines (HAMS; Parr and Russell 1997), which were an influential early suggestion for hierarchical RL. In their original form, HAMs can be seen as imposing potentially very strong constraints on policies, thus making good ones easier to find. However, it is often the case that what makes these constraints appropriate is a form of potential compositionality in the (perhaps approximately) optimal policy; it is just this form of compositionality that our model exploited. HAMs do not have to involve exclusively egocentric information – nevertheless, as we have seen, this can have great benefits in terms of generalisation.

In neural terms, there is some suggestive evidence that the lateral entorhinal cortex (LEC), which has attracted relatively little attention in work on spatial processing in the hippocampal formation, might be involved in the representation of associations in an egocentric or object-centred reference frame. The MEC and LEC are both part of the same structure, but are each the respective termination points of the two highly distinct main input streams into the hipppocampus, known as the dorsal (“where”) and ventral (“what”) pathways (Manns and Eichenbaum 2006). The dorsal stream culminates in the parahippocampal (PHC) cortex and then the MEC (before entering the HPC), while the ventral stream ends in the perirhinal cortex (PRC) and then the LEC. There are PHC/LEC and PRC/MEC interconnections but the majority of information flow seems to run in parallel (Knierim, Neunuebel, and Deshmukh 2014). The suggestion the LEC performs somewhat similar computations to the MEC, but in a different state space, would be parsimonious and might help us further understand hippocampal representational dynamics. In particular, computational similarities between LEC and MEC would allow the plethora of theoretical approaches to understanding the function of latter to illuminate the former.

Of course grid cells in the MEC are easily detectable because of the regularity of their firing fields in a transparently obvious spatial reference frame. Our analysis of the egocentric SRs highlights the difficulty of characterising egocentric associative representations by looking at allocentric “rate” maps. Worse, the coordinate systems in which the egocentric SRs are transparent might have direction dependence and localisation as key components. Some of the issues that arise include the unidirectionality of temporal travel in comparison to spatial travel, as well as the discrete and irregular nature of egocentric states in comparison to allocentric space; this is partly due to the differing saliencies of objects.

There is some evidence that the LEC codes for external objects in an egocentric reference frame (Wang et al. 2018) – we recreated their rate-maps with our model using the agent’s egocentric SR in Figure S1. It is also known that input from the LEC can support hippocampal spatial selectivity (Hales et al. 2014) – as cells with place fields still form in the hippocampus after an MEC lesion – just with decreased spatial precision and decreased long-term spatial stability. A more speculative egocentric correlate is the LEC’s role in associative learning, such as its involvement in recognising associative combinations of objects and contexts (Wilson et al. 2013) as well as its encoding of learned CS-US temporal associations (Suter, Weiss, and Disterhoft 2019) and item-outcome associations (Jun et al. 2024), where LEC neurons have been shown to group items based on their outcomes. The LEC has also been shown to display reward related egocentric encoding (Issa et al. 2024), with distinct populations of neurons representing reward approach, consumption and departure. While the evidence that the LEC represents egocentric relational structure over relations between objects is tentative, the above model principles still apply – especially as an egocentric SR would, by the grouping of different egocentric states across areas of an environment, could also represent and relate objects.

Evidence for related egocentric coding in the HPC includes findings that hippocampal CA1 neurons in bats show angular tuning to goal direction (Sarel and Ulanovsky 2017). By contrast, spatial view cells in the primate hippocampus (Feigenbaum and Rolls 1991) are not truly egocentric as they preferentially code for the allocentric location of space being looked at.

Various facets of the hippocampus and the hippocampal formation may be illuminated by considering a combination of allocentric and egocentric information. One is the concurrent use of spatial and object reference frames in attention (Cave and Bichot 1999; Chen 2012), which is key for the encoding and retrieval of explicit memories from the hippocampus (Muzzio, Kentros, and Kandel 2009). A second is changes in the hippocampal representation of space associated with alterations in the surrounding environment: for instance, hippocampal drift (Ziv et al. 2013) and global (Muller and Kubie 1987), rate (Leutgeb et al. 2005), and partial (Anderson and Jeffery 2003) remapping could be products of different amounts of change in egocentric versus allocentric structure.

In order to focus on the nature of the spatial representation, we considered a very simple problem and domain, with an environment that was deliberately designed to include shared local structure. Navigation in richer environments with more and different obstacles could benefit from non-linear functions of allocentric coordinates and egocentric input, as would arise from multi-layer neural networks. Extending the R_*d*_ function that aligns the reference frames to handle different relationships would could be complex, for instance, with object-based reference frames requiring the alignment of rather different temporal scales of action. It would be interesting to extend our account beyond spatial navigation and explore the benefits in more general problems, where the equivalent of allocentric and egocentric representations could be absolute and relative coding schemes.

In sum, we have offered a perspective on egocentric contributions to spatial behaviour. This suggests that one function of the LEC might be to represent associative egocentric structure; a view that unifies disparate results on its role in representing events (Tsao et al. 2018) and in linking objects and outcomes. Our work suggests that the LEC might play a similar role to the MEC in cognitive mapping, but in a different reference frame. It also shows how cognitive maps might represent different relational structures and how they could be combined to realize flexible cognition.

## Acknowledgments

We thank Jacob Bakermans, Daniel Dombeck, Mathias Sablé-Meyer and Eleanor Spens for their helpful feedback on the manuscript, and Tim Behrens for his support during the project. Funding was from the Max Planck Society and the Humboldt Foundation. PD is a member of the Machine Learning Cluster of Excellence, EXC number 2064/1 – Project number 39072764 and of the Else Krö ner Medical Scientist Kolleg ‘ClinbrAIn: Artificial Intelligence for Clinical Brain Research’.

