## Supplementary Material for "Composing egocentric and allocentric maps for flexible navigation"

### Supplementary Figures

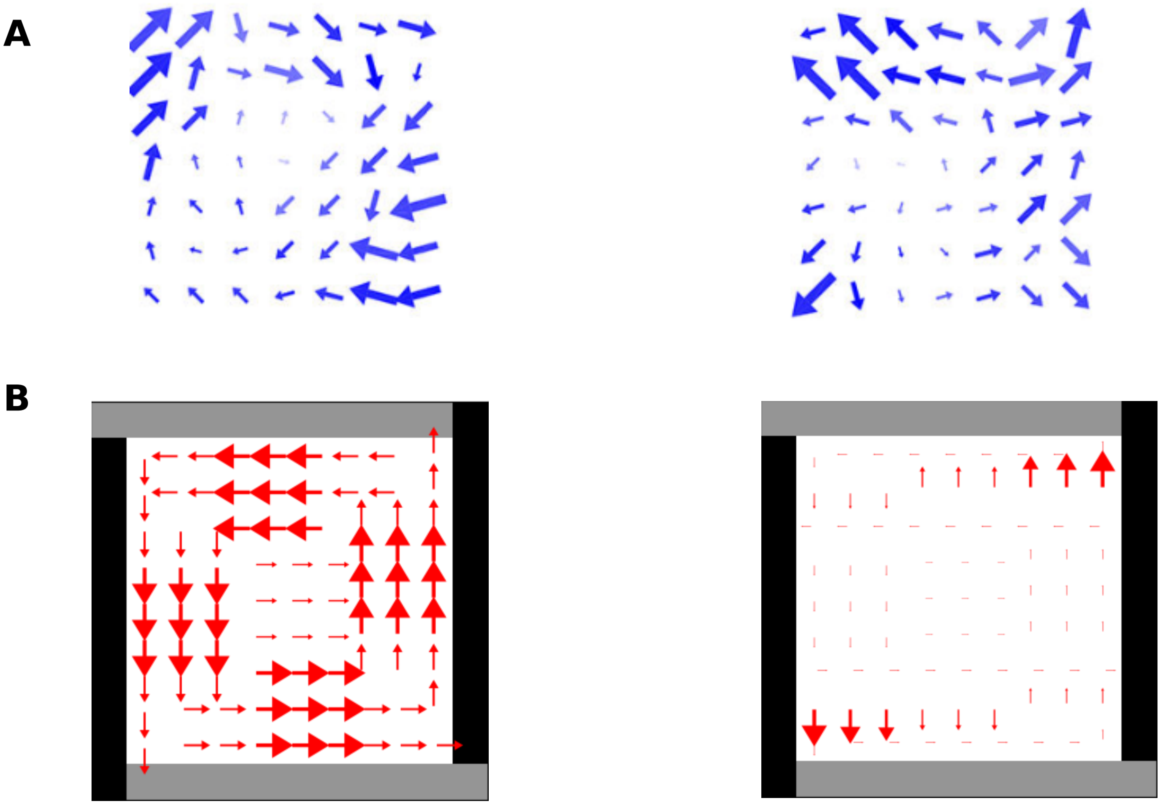

Figure S1: **Tuning field replication.** We create with our model head direction tuning fields that are related to those recorded from the LEC by Wang et al. 2018. LEC rate maps from Wang et al. 2018. **(**A) Two LEC cells. Each plot is a head direction tuning field plot showing preferred head direction (arrow direction) at different locations. Arrow size denotes firing rate. **(**B) Rate maps from our model. Each plot represents an egocentric SR for a new empty environment with two wall colours. We view each state-state egocentric SR as a neuron and plot its head-direction tuning field as follows. We first, for each $(x, y)$ location in the environment, compare the SR's values for the four egocentric states corresponding to different head directions at that location, discounting by total value across all SRs (to overcome the aliasing effects from which our model suffers). Of the four egocentric states SR values, we choose the maximum, and plot an arrow in the corresponding head-direction with magnitude corresponding to the associated SR value. Our model can only capture some aspects of the LEC rate maps. In particular, as there are only four possible head-directions in our paradigm, we cannot capture the diagonal tuning found at some locations. However, our model also finds "neurons" with corner preference and rotational effects, as seen in the LEC neurons.

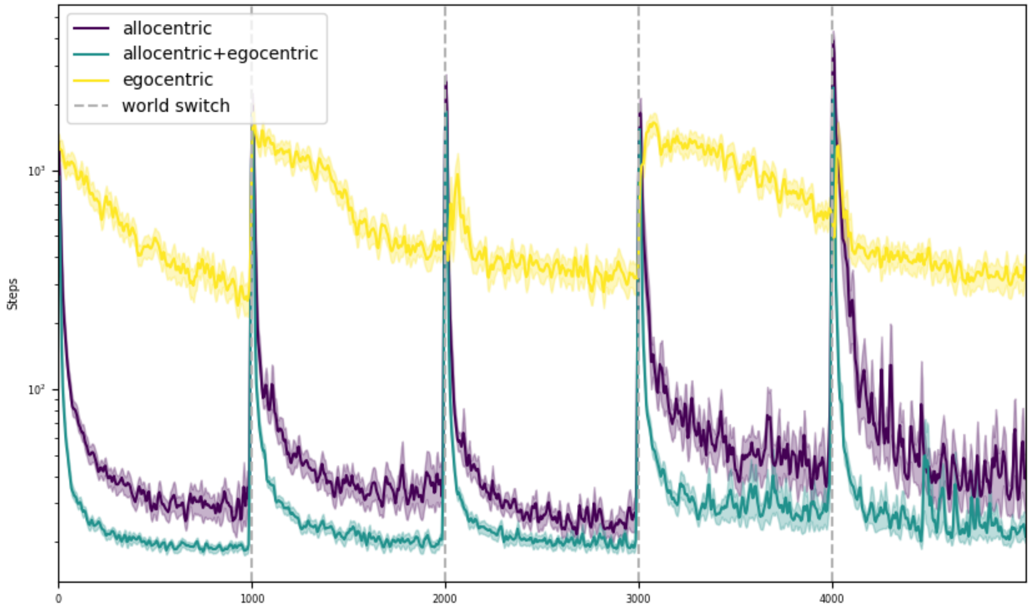


Figure S2: **Full lesion comparison.** Performance of the allocentric+egocentric, allocentric-only, and egocentric-only agents in the standard task paradigm. The egocentric-only agent performs poorly since its representation is local and does not contain sufficient information to find the reward.

| | |
|---|---|
| $\gamma$ | 0.98 |
| $\gamma_E$ | 0.98 |
| $\gamma_A$ | 0.94 |
| $\alpha_E$ | 0.035 |
| $\alpha_A$ | 0.0078125 |
| $\tau$ | 1/100 |
| $\epsilon$ | 0.01 |

Table S1: Model hyperparameters

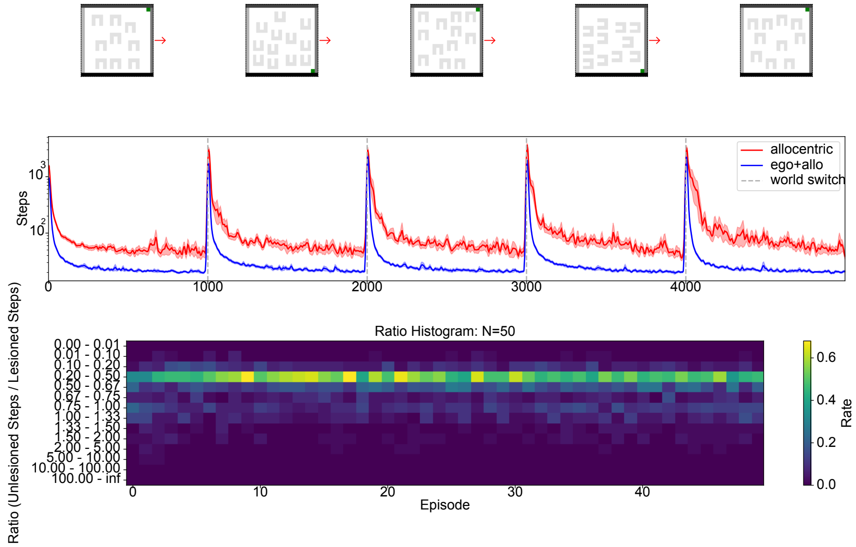


Figure S3: **Task: Aligned, same-size** Upper: performance of the full and lesioned agents across 50 tasks each consisting of 5 environments consisting of identical, randomly positioned barriers which are all aligned to obstruct reward access. Lower: histogram of ratios of number of steps taken per episode of unlesioned to lesioned agents.

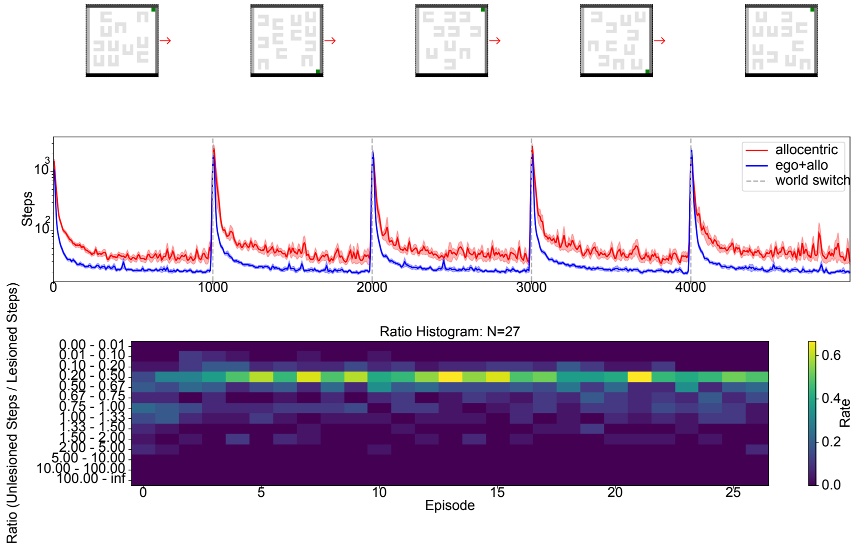


Figure S4: **Task: Unaligned, same-size** Upper: performance of the full and lesioned agents cross 27 tasks each consisting of 5 environments consisting of identical, randomly positioned and oriented barriers. Lower: histogram of ratios of number of steps taken per episode of unlesioned to lesioned agents.

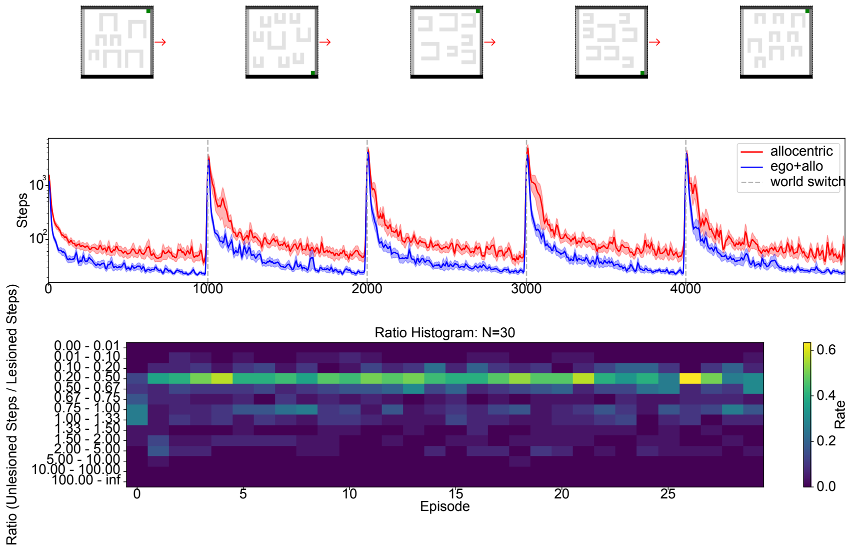


Figure S5: **Task: Aligned, different-size** Upper: performance of the full and lesioned agents across 30 tasks each consisting of 5 environments consisting of randomly sized and positioned barriers, which are all aligned to obstruct reward access. Lower: histogram of ratios of number of steps taken per episode of unlesioned to lesioned agents.

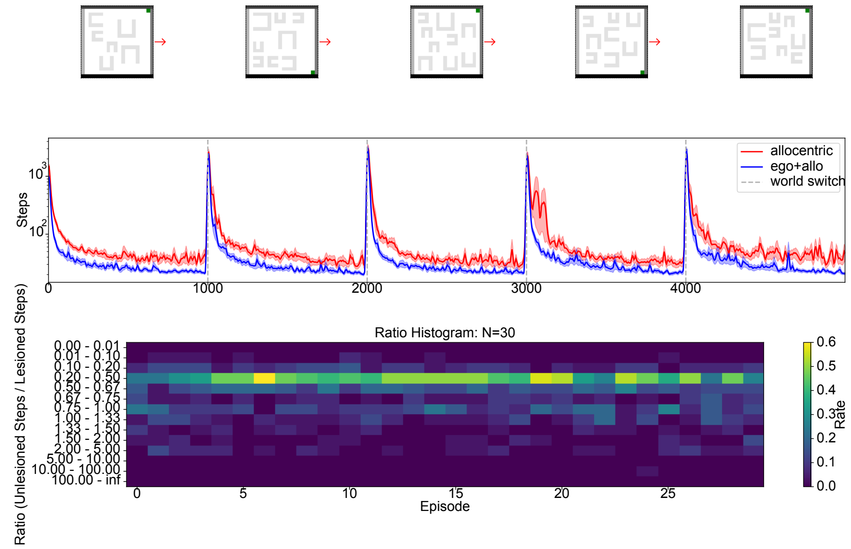


Figure S6: **Task: Unaligned, different-size** Upper: performance of the full and lesioned agents across 30 tasks each consisting of 5 environments consisting of randomly sized, positioned, and aligned barriers. Lower: histogram of ratios of number of steps taken per episode of unlesioned to lesioned agents.

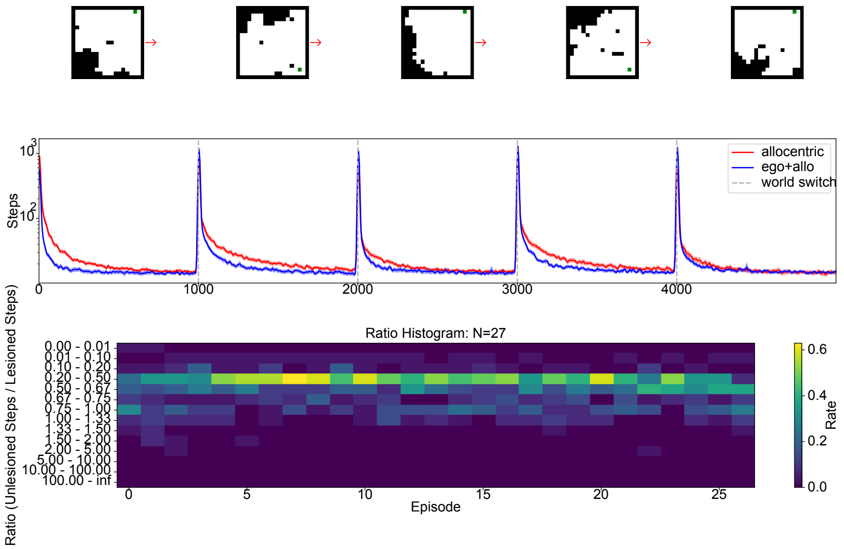


Figure S7: **Task: Randomly generated world** Upper: performance of the full and lesioned agents across 27 tasks each consisting of 5 environments consisting of randomly generated worlds with no shared egocentric structure across environments. Lower: histogram of ratios of number of steps taken per episode of unlesioned to lesioned agents.
